# Velocity-based optoretinography for clinical applications

**DOI:** 10.1101/2022.05.10.491416

**Authors:** Kari V. Vienola, Denise Valente, Robert J. Zawadzki, Ravi S. Jonnal

## Abstract

Optoretinography (ORG) is an emerging tool for testing neural function in the retina. Unlike existing methods, it is noninvasive, objective, and provides information about retinal structure and function at once. As such, it has great potential to transform ophthalmic care and clinical trials of novel therapeutics designed to restore or preserve visual function. Recent efforts have demonstrated the feasibility of the ORG using state-of-the-art optical coherence tomography systems. These methods measure the stimulus-evoked movement of subcellular features in the retina, using the phase of the reflected light to monitor their position. Here we present an alternative approach that monitors the velocity of these features instead. This conceptual shift has significant implications for the nascent field of optoretinography. By avoiding the need to track specific cells over time, it obviates costly and laborious aspects of the position-based approaches, such as adaptive optics, digital aberration correction, real-time tracking, and three-dimensional segmentation and registration. We used this novel velocity-based approach to measure the photoreceptor ORG responses in three healthy subjects. The resulting responses were reproducible and exhibited dependence on dose and retinal eccentricity. The position of the relevant structures was reconstructed through numerical integration of velocity.

## 1. Introduction

Visual information is gathered in the retina by photoreceptors as they absorb photons and convert their energy into membrane potentials in a process known as phototransduction. The resulting signal propagates through several classes of retinal interneurons before being transmitted to the brain via the optic nerve. An overview of the eye’s anatomy and the location of the retina is shown in Fig. 1(a). Assessment of the visual process and its cellular mechanisms is indispensable for disease assessment in the clinic and studies of the mechanisms of disease and efficacy of therapeutic interventions. To that end, a variety of psychophysical and electrophysiological tests have been used. Some of these, such as eye charts and perimetry, are subjective, in the sense that they require feedback from the patient regarding the visibility of stimuli. Subjective tests can be time consuming and suffer from spurious sources of variance such as attention and learning effects. Others, like the electroretinogram (ERG) [1], permit objective measurement of stimulus-evoked electrical activity, but are moderately invasive, requiring placement of electrodes on the cornea and face.

**Fig. 1.**
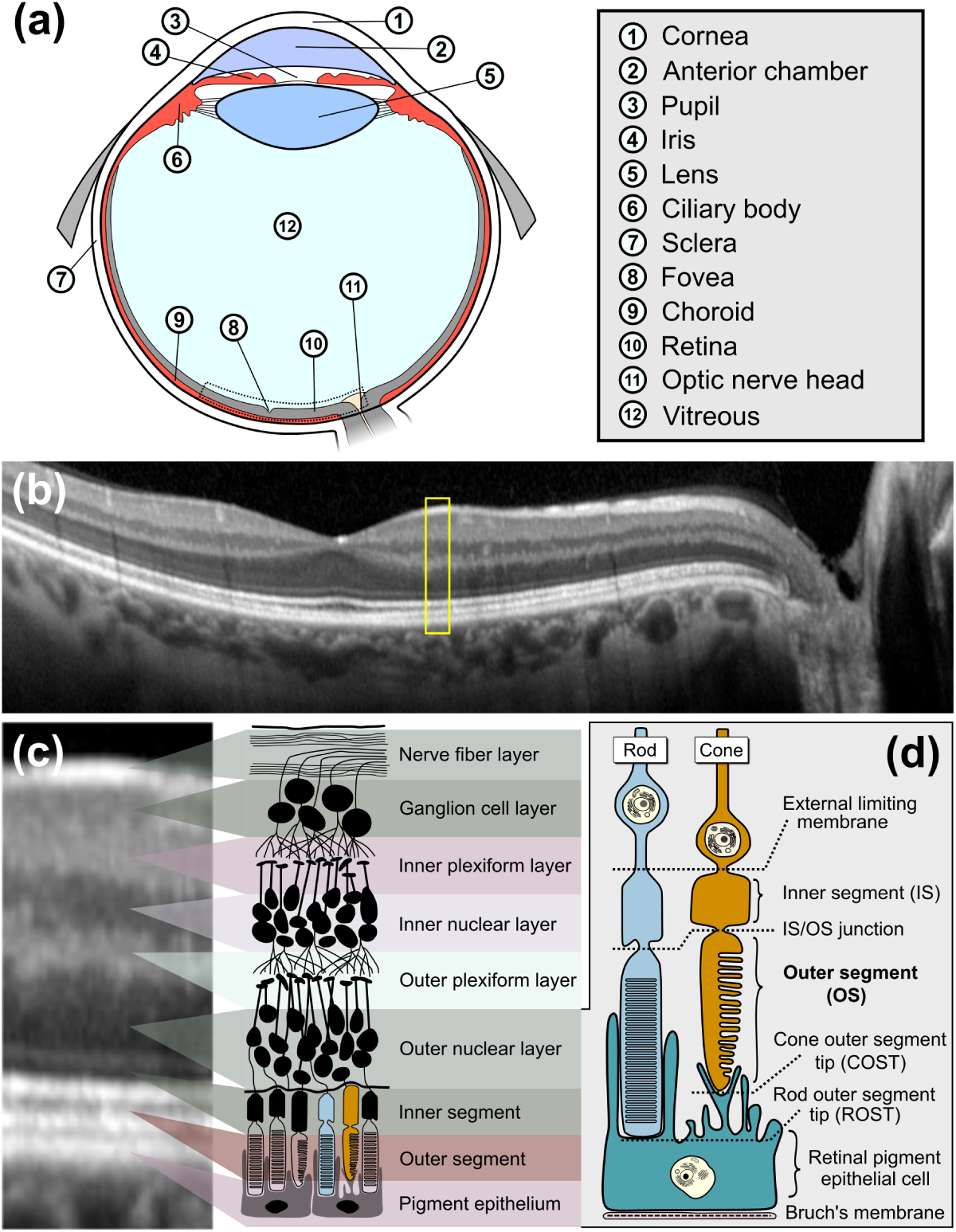
Outer segment elongation can be observed with phase-sensitive OCT. (a) Schematic of an eye indicating key parts of the eye. (b) A wide field-of-view B-scan of the human retina expanding from temporal to nasal across fovea and optic nerve head. c) Drawing shows the different cellular structures of the retina and what they correspond to in a B-scan. (d) An illustration of the human photoreceptor cells with details of different structures. For optoretinography, the outer segment (OS) elongation is observed.

In addition to assessing visual function, clinicians need to observe how the retinal structure is affected by disease. Historically, this was done with slit lamp exams and fundus photography, but over the past two decades, optical coherence tomography (OCT) has become a standard of ophthalmic care. With OCT, the laminar structure of the retina can be visualized non-invasively in three dimensions (Fig. 1(b)) [2]. Even with much progress in the development of tools for both functional and structural assessment, a need exists for an objective, noninvasive assay of retinal function, ideally capable of simultaneously observing the structure of the retina.

As an interferometric imaging modality, OCT records the amplitude and phase of light reflected by the retina. The familiar OCT image is created using only the signal amplitude, while the phase is disregarded. However, the phase of the signal contains information about the position of scattering objects in the scene, and is thus sensitive to tissue movement, even if it is much smaller than the optical resolution of the imaging system [3]. Sensitivity to the movement of a structure depends on its reflectivity: for the brightest features in the ophthalmic OCT image, movements as small as a few nanometers may be measured. This level of sensitivity requires computational methods to correct for the confounding phase shifts caused by eye motion [4, 5].

Because neurons are known to swell and shrink during signaling, phase-sensitive OCT offers the ability to detect whether these cells are responding to stimuli. Early efforts to observe these stimulus-evoked responses in human cone photoreceptors used adaptive optics (AO), which permitted resolution and tracking of single cells, in conjunction with common path interferometry [6]. Since then, interest in this area has grown, and these responses have been successfully measured in cones using detection schemes such as full-field swept-source OCT with digital aberration correction (DAC) [7], flying-spot AO-OCT [8, 9], and line-scanning OCT and AO-OCT [10]. Recently, responses have also been measured in rods, using a multimodal AO-OCT system [11]. These cutting-edge systems all provide measurements of neural responses at the level of single cells, where characteristics of the response may be studied in detail and where disease-related dysfunction manifests earliest. The most commonly reported optoretinographic method has been to localize the two boundaries of the photoreceptor outer segment–the inner-outer segment junction (IS/OS) and the cone or rod outer segment tip (COST or ROST)–and monitor the difference between the phase of light returning from the two structures (see Fig. 1(c-d)). This novel way to observe photoreceptor responses in the retina has been termed *optoretinography* (ORG) [12, 13], and it has been successfully used to classify cones by spectral properties [9] and to detect cone dysfunction in retinitis pigmentosa patients [14].

To our knowledge, the ORG is the only noninvasive, objective test of neural function in the retina that can simultaneously reveal its structure, making it ideal for ophthalmic care and clinical research. However, the advanced imaging systems used to prove the ORG concept pose some challenges for clinical translation. The systems utilizing AO require expensive components and incur additional costs in personnel due to their optical complexity and need for multiple expert operators. In addition, the data rates of these instruments are in the tens of gigabytes per second which, in conjunction with the required data processing, precludes rapid test results at present. Together, these constraints limit the number of healthy and disease-affected eyes that can be tested, and thus the translational utility of the test.

Here, we present a novel ORG approach using a custom OCT system very similar to those currently employed in the clinic. Unlike the research systems described above, it lacks the ability to resolve and track single cells. Instead, the signal processing pipeline was designed around the assumption that the exact cells being imaged at any given time may move out of the field of view at other times and depended on the correlated behavior of adjacent cells to extract a meaningful signal. Phase changes in the IS/OS and COST layers were measured within a time window short enough to ignore retinal movement < 10 ms, and converted into instantaneous, depth-dependent tissue velocities. The resulting series of velocities is related, by integration, to the underlying contraction and expansion of the cone outer segments. Reconstruction of the tissue position may not be necessary anyway, since previous reports suggest that the derivative of the position signal (i.e., velocity) is an advantageous way to quantify it [10, 11].

## 2. Methods

### 2.1. Optical coherence tomography system

The OCT system is illustrated in Fig. 2(a). It used a 1060 nm swept-source (SSOCT; Axsun; Billerica, MA, USA), with a 100 kHz A-scan rate and 100 nm bandwidth. A fiber Bragg grating (FBG) at the source output generated a notch in the acquired spectra, used for spectral alignment of scans. Ten percent of the light from the source was directed to the sample arm, where it was collimated (AC080-10-B; Thorlabs; Newton, NJ, USA) before passing through the galvanometric scanners (6210H & 8310K; Cambridge Technology; Bedford, MA, USA) and a demagnifying telescope *f*_L1_ = 100 nm, *f*_L2_ = 75 mm, creating a 1.2 mm diameter beam on the cornea (pupil plane). The last lens (L2) can be translated to correct defocus. Ninety percent of the source light was directed to the reference arm where it propagated through a polarization controller and dispersion-compensating fiber patch cord. The backscattered light from the eye was combined with the reference arm using a 50/50 fiber coupler. The reference arm power was adjusted using an aperture (A_1_ in Fig. 2(a)) and a retroreflector was translated in one dimension to adjust the reference arm length. The optical power for the OCT in the sample arm at the pupil plane was measured to be 1.8 mW.

**Fig. 2.**
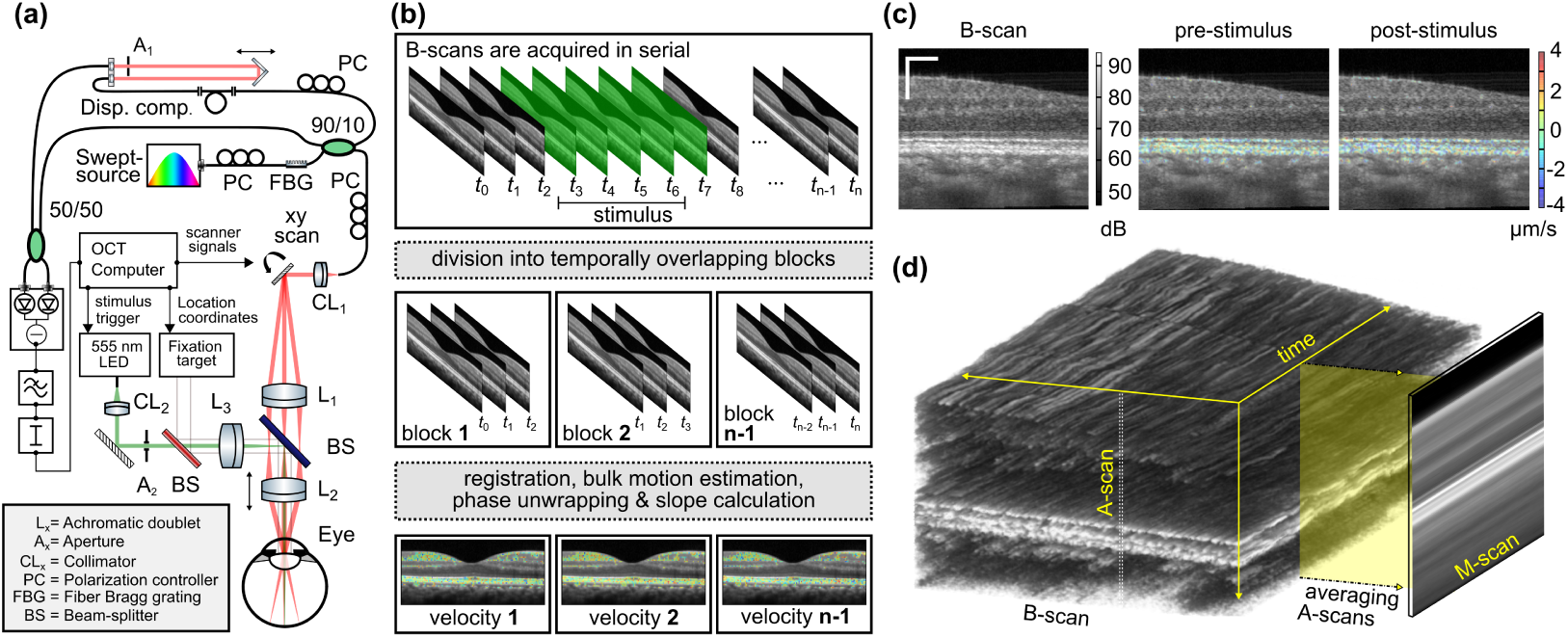
Optical layout and top level data processing pipeline. (a) The SS-OCT system layout (not in scale). (b) A diagram of the ORG data processing. Once the block size width is selected, the phase slopes are obtained from the block data as it moves with a step size of one B-scan through the acquired dataset. c) These phase slopes are calculated from the block for each individual pixel in OCT and visualized after bulk motion filtering. The residual velocities in the inner retina are due to blood flow. (d) The M-scans are generated by averaging A-scans within each individual B-scan, creating an averaged depth profile over time.

To deliver a temporally controlled stimulus flash to the retina, a fiber-coupled 554 nm light emitting diode (MINTF4; Thorlabs; Newton, NJ, USA) was used. Light exiting the fiber was collimated and filtered using a 23 nm bandpass filter centered at 555 nm (FF01-554/23; Semrock; Lake Forest, IL, USA). The stimulus light was combined with the OCT beam path using a dichroic beamsplitter (T715lp; Chroma; Bellows Falls, VT, USA). The stimulus light passes through a non-magnifying telescope *f*_L3,L2_ = 75 mm before entering the eye. The diameter of the stimulus was approximately 360 µm on the retina. The 555 nm stimulus wavelength was selected because it isomerizes L- and M-photopigments equally, which make up the overwhelming majority of cones [15].

The voltage signal from the balanced detector (PDB481C-AC; Thorlabs; Newton, NJ, USA) was filtered with a 150 MHz low pass filter (SLP-150+; Mini-Circuits; Brooklyn, NY, USA) and attenuated 15 dB (VAT-15+; Mini-Circuits; Brooklyn, NY, USA) before being sampled at 12 bits with a ±400 mV range using a digitizer (ATS9350; AlazarTech; Pointe-Claire, QC, Canada). The swept-source provided timing signals from its k-clock and sweep trigger to the digitizer. The analog waveforms to control the galvanometric scanners were generated using a multifunction data acquisition card (NI6251; National Instruments; Austin, TX, USA), and the OCT data acquisition was synchronized with the waveform generation.

The OCT software was developed in C++ and CUDA and has been previously published [16]. When the OCT acquisition started, a 5 V trigger pulse was sent to a function generator (DG4202; RIGOL; Suzhou, JS, China), which then sent the preconfigured stimulus signal to the LED controller (DC4100; Thorlabs; Newton, NJ, USA). By setting the delay, width, and the amplitude of this signal, the stimulus flash could be modified to bleach the desired percentage of photopigment.

### 2.2. Calculating photopigment bleaching

Photopigment bleaching can be calculated using the equation [17]

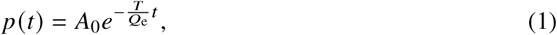

where *A*_0_ is the initial amount of the photopigment, *p* (*t*) is the fraction of the remaining photopigment, *T* is the conventional illuminance of the stimulus flash in Trolands (Td), *t* is the duration of the stimulus (in seconds) and *Q*_e_ is the conventional luminous exposure (in Td ·sec) needed to deplete the photopigment to 1/*e*. For our calculations, we’re using the value of 2.4 × 10^6^ Td sec for *Q*_e_, following Rushton and Henry [17]. The conventional retinal illuminance *T* can be calculated from the luminous power using the following equation [18]:

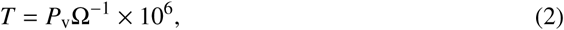

where *P*_v_ is the photopic luminous power in lumens and Ω is the solid angle measured from the nodal point of the eye. To convert our radiometric measurement into photometric we use:

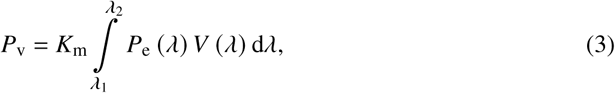

where *K*_m_ is a conversion constant equal to 683 lm W, *P*_e_ is the radiant power and *V* (*λ*) is the photopic luminous efficiency function. For simplicity, we assumed our light to be monochromatic with *λ*_c_ = 555 nm, resulting in *V* (*λ*) of 1. Finally, by combining equations (2) and (3), we get:

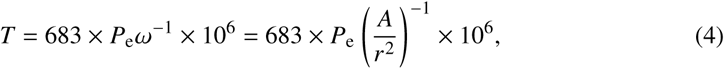

where the unit of *P*_e_ is the radiant power in watts, *A* is the illuminated area, and *r* is the distance from the eye’s nodal point to the retina (which we assumed to be 17 mm). As the stimulus size on the retina has a diameter of 360 µm, we have all the information needed to convert the optical power measured from the pupil plane into Trolands, and then use it in equation (1). The resulting bleaching levels and corresponding optical powers are listed in Table 1.

**Table 1.** Different parameters obtained from photopigment bleaching calculations. All calculations were done with a pulse width of 30 ms. For simplicity, ocular transmission was assumed to be 1.

| Bleaching [%] | Stimulus [μW] | # of photons ($\times 10^{11}$) | Photon density [photons/μm<sup>2</sup>] |
| --- | --- | --- | --- |
| 66 | 45 | 37.7181 | $3.7 \times 10^7$ |
| 33 | 16.5 | 13.8300 | $1.4 \times 10^7$ |
| 17 | 7.7 | 6.4540 | $6.3 \times 10^6$ |
| 8 | 3.45 | 2.8917 | $2.8 \times 10^6$ |
| 4 | 1.7 | 1.4249 | $1.4 \times 10^6$ |

### 2.3. Imaging protocol

All imaging procedures were in accordance with the Declaration of Helsinki and were approved by the UC Davis Institutional Review Board. Written informed consent was obtained from all participants following an explanation of experimental procedures and risks, both verbally and in writing. To ensure safe imaging, the laser safety limits were calculated using the latest standard [19].

The use of mydriatic drops was not necessary. A custom bite bar was fabricated for each subject to position and stabilize their head. For ORG measurements, subjects were dark adapted for five minutes and then asked to look into the system and fixate on the fixation target (Fig. 2(a)). The OCT system was configured to scan in one dimension only, acquiring a series of B-scans in a single location. B-scans consisting of 250 A-scans (scan width approx. 750 µm) were collected at a rate of 400 Hz. B-scans were collected before and after the stimulus flash, with the 40 scans before and after the flash used for subsequent analysis.

Two experimental parameters were explored: distance from the foveal center and stimulus dose. Generally, multiple retinal loci were imaged after each period of dark adaptation. This was done in order to vary the distance from the foveal center or to collect multiple trials at a single distance. In the latter case, the multiple loci lay in concentric, iso-eccentric rings, with measurements taken at up to five loci.

All measurements were done on the temporal side of the retina, at distances of 2°, 4°, 6° and 8° away from the foveal center. A total of three different subjects were used in this study, all having healthy maculae. Images were acquired at each of the four distances from the fovea, in all three subjects. In one of the subjects, the stimulus dose was varied to achieve six different L/M-photopigment bleaching levels, between 0 % and 66 %, with ORG measurements collected at four retinal eccentricities for each bleaching level.

### 2.4. Signal processing

The recorded signal was processed in two stages. First, the raw spectral data were converted into cross-sectional OCT images (B-scans), using well-established approaches in the OCT literature. In short, raw digitized spectra were aligned using the FBG notch to create phase-stable B-scans. After the alignment, DC-bias was estimated and removed. The spectra were then corrected for dispersion mismatch between the two arms using numerical dechirping, and complex-valued B-scans were generated by Fourier transforming the processed spectra. B-scans were flattened, such that the IS/OS and COST reflections lay at the same height for each A-scan in the image. Flattening was done by linear shearing while optimizing the heights of the corresponding peaks in the laterally averaged profile. In the second stage, the ORG signal was extracted. Complex B-scans were converted into estimates of tissue velocity as follows (Fig. 2(b)). A moving, 10 ms time window was used to select groups of five sequential B-scans at a time. A histogram-based bulk-motion correction algorithm [4] was utilized to compensate for the axial eye movement during the 10 ms interval, with motion corrected relative to the first B-scan in the series. After bulk-motion correction, the resulting complex data cube (V) may be described as:

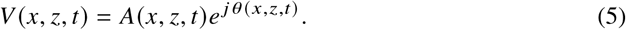

where *x* and *z* are the lateral and depth coordinates, respectively, and *t* is time within the window, i.e. 0 ms ≤ *t* ≤ 10 ms. The phase data cube *θ*(*x, z, t*) was unwrapped in the temporal dimension by adding or subtracting 2*π* to *θ*(*x*_*p*_, *z*_*q*_, *t*_*r*_) in order to minimize *θ*(*x*_*p*_, *z*_*q*_, *t*_*r*_) - *θ*(*x*_*p*_, *z*_*q*_, *t*_*r*-1_). This step was performed for each spatial coordinate pair (*x*_*p*_, *z*_*q*_) in the volume. After unwrapping, a rate of phase change was computed for each coordinate pair by performing a least-squares linear fit with respect to *t*, giving 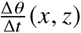 in rad/s. From this, we calculated the instantaneous velocity for each spatial location:

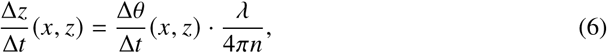

where *λ* = 1060 nm and *n* = 1.38, the nominal refractive index of the eye. An amplitude B-scan, along with pseudocolor overlays of instantaneous pre- and post-stimulus velocities is shown in Fig. 2(c). The next step in quantifying the response was averaging both the B-scan amplitude and 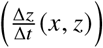 in the lateral dimension, giving instantaneous, depth-dependent measures of backscattering and velocity 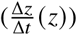, respectively. By shifting the 10 ms window by one B-scan period (2.5 ms) at a time, a time series of depth profiles can be constructed, separately for reflectivity and velocity.

Both of these can be visualized in time-depth coordinates, as M-scans (see Fig. 2(d)). Because the signal-to-noise ratio of the phase changes and velocities are highest when the OCT amplitude is high [3], velocity overlays are only shown for the brighter portions of B-scans and M-scans. From, the velocities of the IS/OS and COST layer movements were extracted, and the From 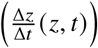, the velocities of the IS/OS and COST layer movements were extracted, and the difference between them is the velocity of contraction/elongation of the OS in the region, which we term *v*_OS_ *t* or simply *v t* hereafter. For each experimental condition, multiple measurements of *v* (*t*) were acquired. The time-varying mean and standard error of the mean (SEM) were calculated from these, and used to generate the plots in Fig. 4.

Preliminary examination of the data revealed a biphasic response (as shown in Fig. 3(d)): an initial OS contraction *v* (*t*) < 0 over the first 10 ms, followed by an elongation *v* (*t*) > 0 that appeared to be somewhat stable between 20 and 40 ms. Motivated by these features, as well as our previous scanning ORG measurements [8, 11], we devised several parameters to quantify the responses (illustrated in Fig. 3(d)): the most negative velocity after stimulus *v*_min_, the greatest positive acceleration *a*_max_, and the time-averaged velocity between 20 and 40 ms after stimulus 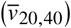. A virtue of these figures of merit is that they depend on data collected within 40 ms of the stimulus flash, thereby avoiding corruption by reflexes caused by the stimulus, such as blinks and saccades. For all measurements, the OS length was recorded as well.

**Fig. 3.**
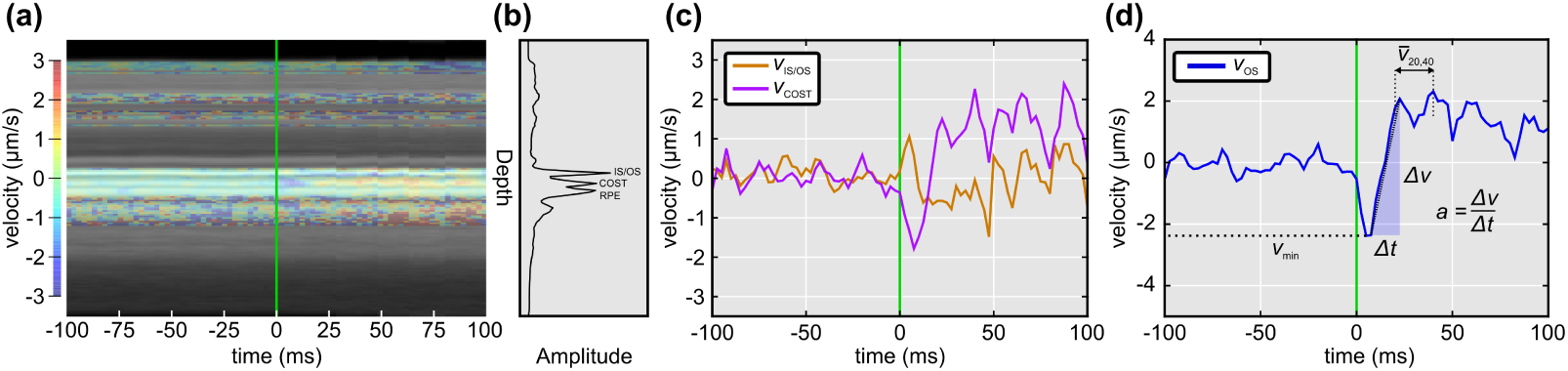
The extraction of velocity from the M-scan. (a) M-scan of the dataset where serial B-scans were acquired. The overlaid color-coding represents the pixel-level velocity values based on phase changes, and the green line indicates stimulus onset. (b) Axial depth profile produced by averaging the M-scan over time. The important peaks here are the inner segment/outer segment junction (IS/OS) and cone outer segment tips (COST) that are used for analysis. (c) Velocity plotted as a function of time for two different layers, IS/OS and COST. Both show rapid changes occurring after the stimulus flash is applied at 0 ms. (d) By subtracting the velocity responses from c), we can then plot the outer segment elongation velocity curve. After the stimulus at 0 ms. the velocity of elongation rapidly changes; an initial contractile stage is followed by a longer period of elongation after ~20 ms. Panel (d) illustrates three approaches we explored for quantification of the OS response: the minimum (i.e., most negative) velocity *v*_*min*_, the average velocity between 20 to 40 ms 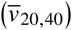, and the maximum acceleration *a*_*max*_.

**Fig. 4.**
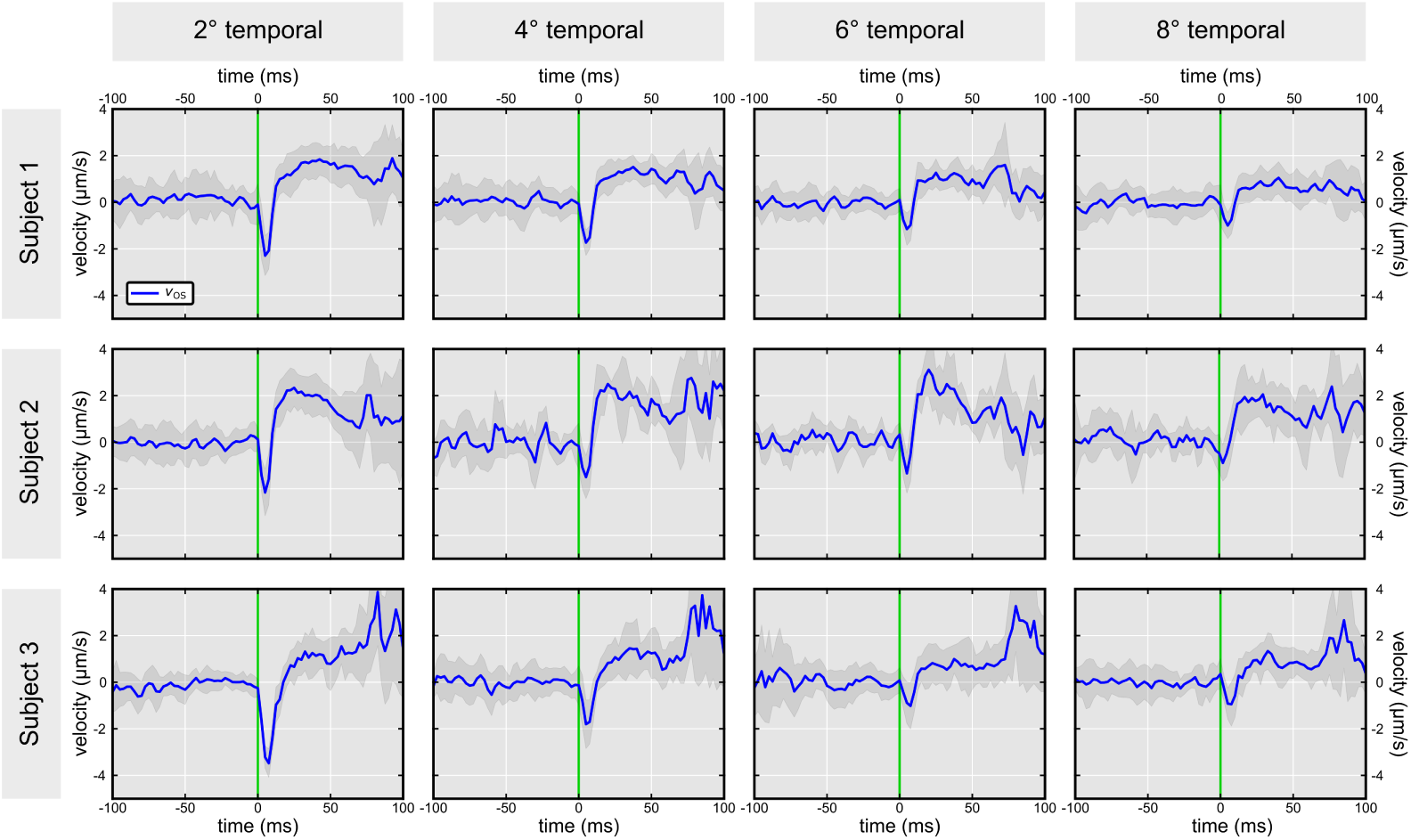
Optoretinograms from three different subjects at four different locations in the temporal retina. Stimulus flash onset was at 0 ms. The blue line shows the average response over five trials, with the shaded gray area indicating one SEM. Retinal eccentricity of the measurement (in visual angle from the foveal center) is indicated in the bottom right of each plot. Two features were evident in all of the measurements: an initial negative velocity (approximately 0 to 10 ms), followed by a longer period of positive velocity (approximately 10 to >50 ms). The magnitude of both features appeared to fall with increasing retinal eccentricity. Responses appeared to become noisy after approximately 70 ms, possibly indicating the presence of reflexive eye movements.

To compare the present measurements of OS velocity to earlier ORG measurements, we reconstructed the OS length response by numerical integration of velocity.

## 3. Results

Figure 3 illustrates an individual ORG measurement. The M-scan (Fig. 3(a)) shows the amplitude of back-scattered light as a function of time and depth in grayscale with tissue velocity superimposed in pseudocolor. The amplitude is plotted as a function of depth in Fig. 3(b), where the peaks originating from the IS/OS and COST are visible. The corresponding tissue velocities can be observed to change in Fig. 3(a), following onset of the stimulus flash (green line). The velocity changes seen in the inner retina and choroid coincide with vascular plexuses, and are thus presumably due to blood flow. The changes due to cone photoreceptor responses can be seen in the IS/OS and COST layers of the M-scan, especially when these velocities are plotted (Fig. 3(c)).

In the first five milliseconds after bleaching, the IS/OS velocity becomes positive (corresponding to downward movement, in the M-scan) while the COST velocity becomes negative (corresponding to upward movement). The movement of these features toward one another is consistent with contraction of the OS. Over the subsequent ten milliseconds, the velocities of the two layers reverse, with the IS/OS moving upward and the COST moving downward, consistent with OS elongation. When the velocity of these two layers is subtracted, the end result is the rate of OS length change (*v*_OS_) as seen in Fig. 3(d). The observed results are consistent with previous reports using adaptive optics, which showed that the OS contracts initially (< 10 ms after stimulation), and elongates after that for >100 milliseconds. Proposed figures of merit for quantifying ORG responses are illustrated in Fig. 3(d).

Figure 4 shows the velocity responses measured at different eccentricities in three different healthy subjects. Each blue line is an average of between five and ten trials, with the gray area representing the standard error of the mean (SEM). A response was visible in all individual trials (as illustrated in Fig. 4), and can be seen in the average response across all subjects and eccentricities. Both the contractile and elongation periods appear to scale with the eccentricity, reducing toward the periphery in all three subjects. In many individual trials, velocity measurements became noisy around 70 ms after the stimulus flash, and this is evident in the higher SEM visible in that portion of several of the plots. We speculate that the noise is a consequence of a reflexive eye movement that reduces the effectiveness of our current bulk-motion algorithm. Since the following statistical analysis utilized B-scans collected within 40 ms of the stimulus onset, they were unaffected by this noise.

The eccentricity-dependence of the response is visualized in Figs. 5(a) and 5(b). Here, parameters *v*_min_ and *a*_max_ are used to illustrate the trends. The most negative velocity *v*_min_, which corresponds to the largest rate of contraction, decreased in magnitude between 2° and 6°,with similar values at 6° an d 8°. The maximum acceleration of OS elongation, (*a*_max_), decreased in magnitude between 2° 1.5 to 2.1 µm/s^2^ and 6° 1.1 to 1.5 µm/s^2^. The velocities are also expressed as fractions of the OS length per unit time (right y-axis, plotted in gray), which flattens the dependence of both aspects of the response on eccentricity.

**Fig. 5.**
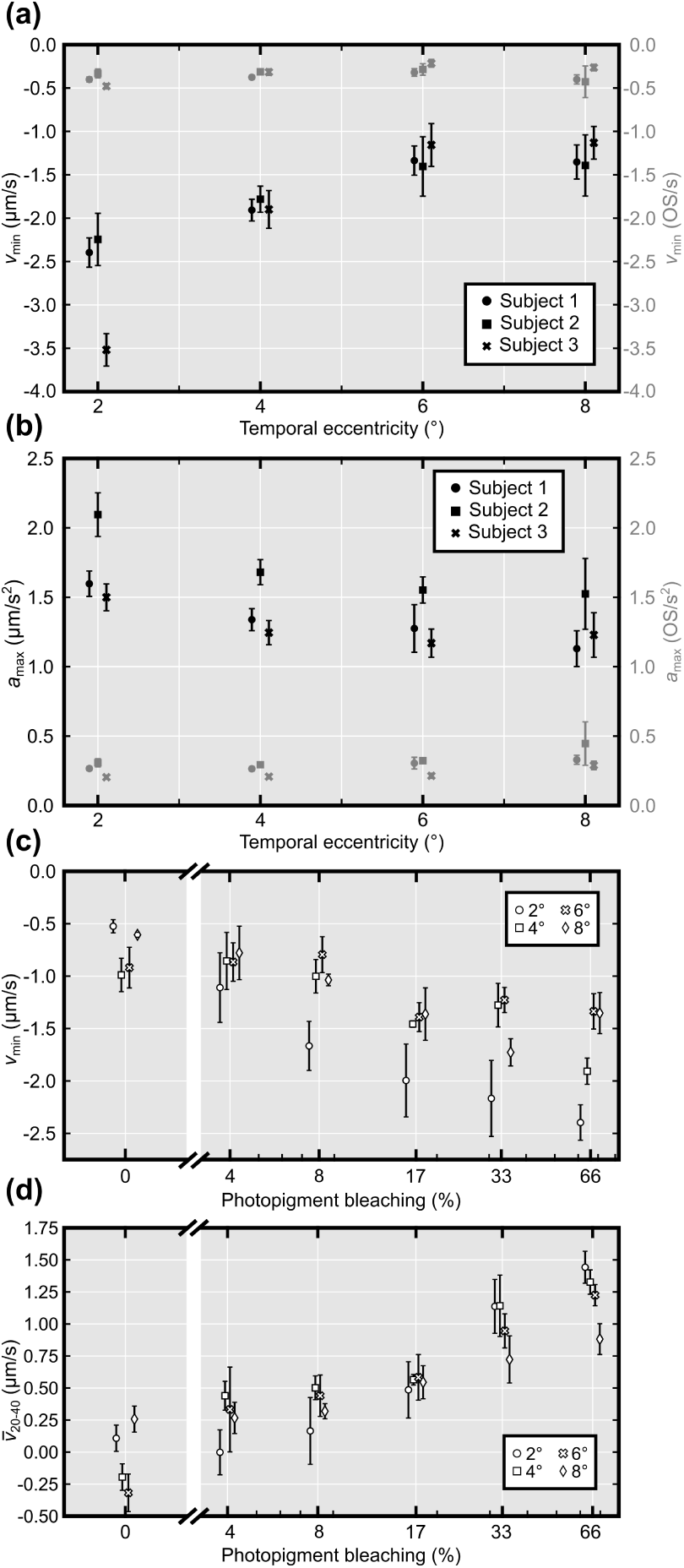
ORG data plotted as a function of eccentricity (a,b) and pigment bleaching (c,d). The most negative velocity (*v*_min_), observed within 20 ms of stimulus onset, reduced in absolute value with increasing visual angle; the fastest OS contractions were observed closest to the foveal center, where the cone OS is longest, as shown by the black markers in (a). The maximum OS acceleration (*a*_max_) was similarly highest nearest the fovea, as shown by the black markers in (b). When these two values were visualized as fractions of OS length instead of physical length, both relationships appeared to be substantially flattened, as shown by the gray markers in (a) and (b). Both contraction velocity (*v*_min_) and average elongation velocity between 20 to 40 ms *v*_20,40_ appeared to depend on the stimulus dose, having the largest magnitudes in response to the brightest stimuli, as shown in (c) and (d), respectively. In panel (c), the unexpected non-zero contractile velocity in the absence of stimulus may be due to bias introduced by the quantitative parameter *v*_min_, which will be negatively biased due to noise.

Dose-dependence of the response is visualized in Figs. 5(c) and 5(d), using log scale for the x-axis (bleaching). Figure 5(c) illustrates the dependence, at all eccentricities, of the contractile response on dose, with higher doses having the largest (most negative) velocities. Contractile velocity was very similar at doses of 0 % and 4 % but they appear to be different in the case of elongation velocity. When the elongation was quantified as the average velocity between 20 and 40 milliseconds post-stimulus 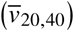, a similar trend was visible, with brighter stimuli resulting in faster elongation. In response to the brightest stimuli (66 % bleaching), elongation velocities were in the range of 1.00 to 1.25 µm s. Qualitatively, dose and response appear to have a log-linear relationship between 4 % and 66 % bleaching. In all panels of Fig. 5, data points were offset horizontally for readability.

Numerical integration of the OS velocity responses is shown in Fig. 6. The velocity of the rapid elongation stage, between approximately 10 and 100 ms, appears to be about 1 µm s. This substantially lower than previous studies, which reported a velocity of approximately 3 µm s to a 70 % bleach [8] and velocities approaching 3.9 µm s and 4.6 µm s for similar flashes [10]. We note that numerical integration is not necessary for this comparison, as the slope of the integrated curve is the same as the time-averaged velocity in the same 10 to 100 ms interval.

**Fig. 6.**
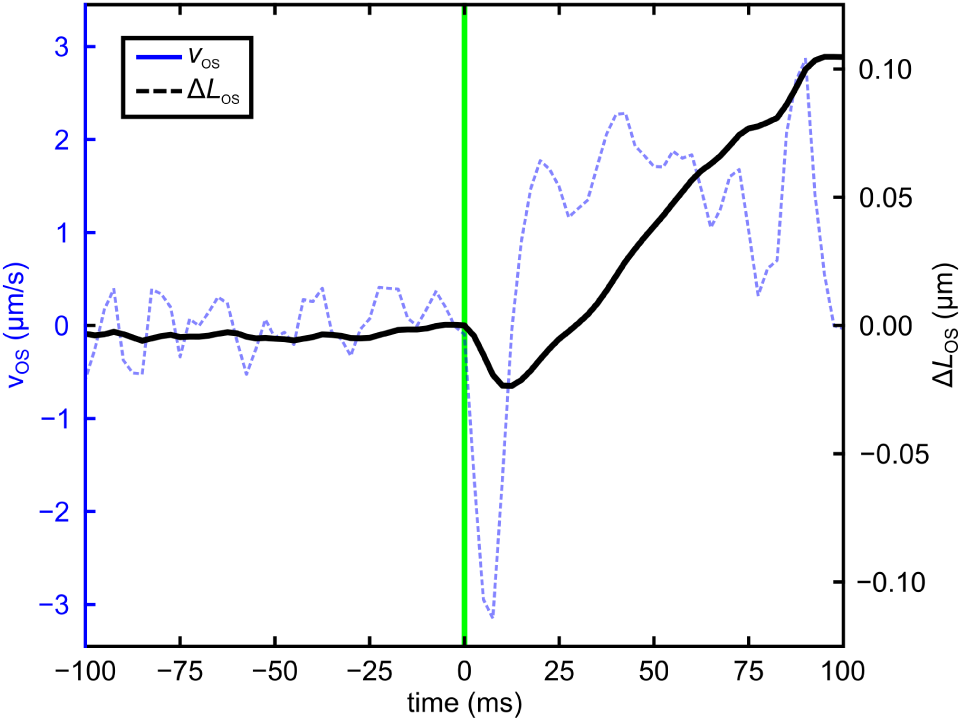
Outer segment velocity data numerically integrated to reconstruct the length-based ORG response. The blue line plotted on the left axis shows an example of a response acquired at 2° temporal in Subject 1, to a stimulus flash that bleached 66 % of L- and M-photopigment. The dashed black line plotted on the right axis is the numerical integration from 100 to 100 ms. The slope of the resulting estimate of elongation from 20 to 100 ms is approximately 1 µm s, lower than values reported by others using position-based methods.

## 4. Discussion

Optoretinography offers a completely new tool for ophthalmologists and vision scientists to produce meaningful and quantifiable data about human vision. The stimulus-evoked contraction and elongation of the OS has now been independently reproduced by several research groups using hardware adaptive optics and/or full-field OCT with digital aberration correction (DAC) to visualize individual photoreceptors. The conventional (non-AO) optoretinographic method demonstrated here depends on the correlated movement of neighboring tissue, and thus circumvents the need for cellular resolution and tracking. Compared to more advanced techniques, this method offers a simpler imaging system, reduced data sizes, faster signal processing, and smaller demands on personnel. One of its key advantages is potentially straightforward incorporation into existing commercial technology. The data shown in Figs. 5a and 5b required approximately ten minutes per subject to acquire, including dark adaptation, and required just one operator. Our data processing software was written in a high-level language (Python/Numpy) and is thus not optimized for real-time use. Nevertheless, results are available within minutes, making the approach attractive to clinicians and large-scale studies. As shown in Fig. 6, the main result of this method–a time series of OS velocity–can be integrated to approximate the results of earlier, position-based optoretinographic methods. Congruity between the two results is an advantage to both, as they can contribute to the same growing body of literature and data. The present results reveal velocities that are lower by a factor of two to three than earlier position-based results. The pattern of contraction and elongation demonstrated here bears significant similarity to earlier results, including the timing of the early and late stages. This suggests that the biophysical processes underlying the responses are the same. The reason for the discrepancy in velocities is not known, but could potentially be due to optical factors such as the role played by the lateral PSF size and contributions of portions of the tissue that might be stationary. We are optimistic that such systematic differences between the approaches can be resolved by calibration.

The clear relationship between ORG responses and distance from the fovea (Figs. 4, 5(a), and 5(b)) may be due to differences in outer segment anatomy, and normalizing by OS length seems to weaken this relationship. When designing the imaging protocol, we strove to avoid realignment of the optical system with changes in fixation. Given that, and the fact that this method does not require dilation of the pupil, there is a risk that the subject’s iris could clip the 4 mm diameter stimulus beam after changing fixation. We did not actively monitor this, but it could be done with a real-time pupil camera programmed to detect iridial reflexes. Alternatively, a brighter stimulus source could be used, permitting the beam diameter at the pupil to be smaller. If the OCT and stimulus beam diameters were the same, the OCT signal itself could be used to calculate the stimulus efficiency (not withstanding chromatic differences). Moreover, because the trends were similar among subjects, regardless of which fixation target location was used for initial alignment, we do not believe that the stimulus beam was clipped.

A distinct trend was also observed when we altered the stimulus energy (Figs. 5(c) and 5(d)). Photopigment bleaching is dependent on stimulus dosage between 8 % and 66 % pigment bleaching values. Establishing dose-dependence is a critical step in developing novel functional assays. The dose-dependence observed here are consistent with similar observations of dose-dependence using position-based optoretinographic methods, providing further assurance of the interoperability of these two methods.

We observed that optoretinograms were reliably produced from all tested eccentricities when the stimulus energy bleached more than 8 % of the photopigment. While this sensitivity is lower than what has been reported with AO-OCT systems, the noise floor of the present method could potentially be reduced with improved bulk-motion correction. The 30 ms duration of the stimulus flash was required to reach a photopigment bleaching rate of 66 %, due to limitations in the source power and our optical design. A more powerful source and redesign of the stimulus channel could permit shorter, more intense flashes, which would likely improve the method’s sensitivity. The odd result in Fig. 5(c), that *v*_min_ was non-zero even in the absence of a stimulus flash (0 % bleaching) may be a consequence of noise combined with the bias involved in identifying the minimum (most negative) velocity. This bias probably limits the sensitivity of this figure of merit. One of the motivating assumptions of this work is that there are finite time intervals over which the retina is effectively motionless, i.e., that the axial component of motion is small with respect to the wavelength of the light source and that the lateral component is small with respect to the diffraction-limited spot size (or speckle size). While this assumption appeared largely to be true, evidence of motion artifacts were present in many of the measurements. A potentially powerful way to improve the sensitivity of this approach would be to detect and filter movement artifacts. When analyzing a series of B-scans to measure instantaneous velocity, we monitored the residual least-square error and the correlation of B-scan amplitude. Preliminary investigation showed that both estimates of error were correlated with deviations from expected velocity measurements.

Our results reveal a biphasic ORG response, consisting of an initial contraction and later elongation of the OS. These stages of the response are thought to have different physiological origins, and may thus confer distinct clinical utilities. Zhang et al., by demonstrating a lack of elongation in mutant mice with a dysfunctional G-protein transducin, suggested that transducin subunit dissociation might drive ORG elongation osmotically [20]. Pandiyan et al. hypothesized that the contractile response seen in the ORG is due to the early receptor potential (ERP) [10].

For optoretinograms presented here we used a simple OCT B-scan flattening procedure. In future studies, a segmentation-based approach is needed. Adaptive optics studies of photoreceptor morphology [21, 22] have revealed axially staggered locations of IS/OS and COST in neighboring cones. Moreover, the B-scans shown above illustrate that the ideal depth at which to measure the phase of these surfaces can change from A-scan to A-scan. Instead of computing the phase at the same depth for all A-scans, segmentation would permit measurement of phase at the most salient (and brightest) depths, and thereby improve the ORG sensitivity. It would also allow the investigation of eccentricities closer to the fovea, where the thickness of retinal layers can vary substantially within small visual angles. It could potentially help in the imaging of disease-affected retinae as well, where the layers may be sporadically deformed by edema, drusen, or degeneration.

Lastly, the use of a finite temporal window to calculate velocity is equivalent to convolution of the underlying signal with a sinc function, which limits temporal bandwidth and causes underestimation of instantaneous velocity. Since the convolution kernel is known exactly, potential exists for computational correction of the measurements and estimation of true velocities.

We have shown that it is possible to obtain ORG responses using an SS-OCT system without adaptive optics. The simplicity of the imaging system permits straightforward integration of this method to existing OCT systems. While more research is needed, these preliminary results indicate a promising new approach to produce optoretinograms, paving the way for a clinical ORG system. Future work will include refinement of the optical and computational aspects of the system, further testing of subjects with and without retinal disease, and comparison of the approach to other conventional (non-AO) imaging-based approaches [23, 24].

## Funding

This research was funded by NIH grants R00-EY-026068 (RSJ), R01-EY-033532 (RSJ), R01-EY-031098 (RJZ), RO1-EY-026556 (RJZ) and P30-EY-012576.

## Acknowledgments

The authors would like to thank Dr. Yifan Jian for providing the OCT acquisition software, Prof. John Werner for helping with theoretical calculations and Susan Garcia for clinical assistance.

## Disclosures

The authors declare no conflicts of interest.

## Data availability

Data underlying the results presented in this paper are not publicly available at this time but may be obtained from the authors upon reasonable request.

## Notes

### Competing Interest Statement

The authors have declared no competing interest.

